# Mechanoreceptive Aβ primary afferents discriminate naturalistic social touch inputs at a functionally relevant time scale

**DOI:** 10.1101/2023.07.22.549516

**Authors:** Shan Xu, Steven C. Hauser, Saad S. Nagi, James A. Jablonski, Merat Rezaei, Ewa Jarocka, Andrew G. Marshall, Håkan Olausson, Sarah McIntyre, Gregory J. Gerling

**Affiliations:** University of Virginia; Linkoping University; Umea University; University of Liverpool

## Abstract

Interpersonal touch is an important part of our social and emotional interactions. How these physical, skin-to-skin touch expressions are processed in the peripheral nervous system is not well understood. From single-unit microneurography recordings in humans, we evaluated the capacity of six subtypes of cutaneous afferents to differentiate perceptually distinct social touch expressions. By leveraging conventional statistical analyses and classification analyses using convolutional neural networks and support vector machines, we found that single units of multiple Aβ subtypes, especially slowly adapting type II (SA-II) and fast adapting hair follicle afferents (HFA), can reliably differentiate the skin contact of those expressions at accuracies similar to those perceptually. Rapidly adapting field (Field) afferents exhibit lower accuracies, whereas C-tactile (CT), fast adapting Pacinian corpuscles (FA-II), and muscle spindle (MS) afferents can barely differentiate the expressions, despite responding to the stimuli. We then identified the most informative firing patterns of SA-II and HFA afferents’ spike trains, which indicate that an average duration of 3-4 s of firing provides sufficient discriminative information. Those two subtypes also exhibit robust tolerance to shifts in spike-timing of up to 10 ms. A greater shift in spike-timing, however, drastically compromises an afferent’s discrimination capacity, and can change a firing pattern’s envelope to resemble that of another expression. Altogether, the findings indicate that SA-II and HFA afferents differentiate the skin contact of social touch at time scales relevant for such interactions, which is 1-2 orders of magnitude longer than those relevant for discriminating non-social touch inputs.

## Introduction

Touch is an often used medium for facilitating social relationships and interactions. For example, one might lightly tap another person to get their attention, or stroke a partner’s arm to offer a sense of calm. Between people in close relationships, and even between strangers, many social touch expressions are intuitively understood [1]–[6]. The appreciation of emotion is commonly thought to be a centrally mediated process performed by frontal and temporal brain structures that integrate a multitude of peripheral and cross-cortical sensory information [7]. However, the peripheral nervous system may already be organized to facilitate the selection and processing of potentially socially relevant stimuli [8]. Reliable signaling from peripheral afferents could form the basis of the somatosensory and affective perception in the central nervous system. In our evolutionary history, such peripheral encoding may also have acted as scaffolding for the development of cross-sensory, cortical processing of emotion [9].

Among peripheral tactile afferents, percepts tied to social and emotional touch are thought to be influenced prominently by C-tactile (CT) afferents [10]–[12]. These afferents have been studied mostly in hairy skin, and can be preferentially activated by light stroking contact at 1-10 cm/s velocities [10], [13] and temperatures similar to human skin [14]. Their firing frequencies have been correlated with subjectively perceived pleasantness [10] and this correlation has been widely and reliably reproduced on the population level. Recent work has, however, encountered difficulty in reproducing such trends among individual participants [15], which suggests a more complex view of pleasantness and affective touch and a plausible role of other afferent types. Meanwhile, the firing properties of CT afferents have mainly been characterized in response to controlled stimuli [10], [12], [14], [15], such as rotary actuated brushing. Their firing responses have been explored less under naturalistic, human-to-human touch.

In contrast to CT afferents, low-threshold mechanosensitive (LTM) Aβ afferents have been investigated in a wider variety of scenarios, especially in discriminative touch. Pre-defined, well-controlled mechanical stimuli have been used to decouple and examine stimulus attributes, one at a time [16]–[19]. Across these studies, different Aβ subtypes are dominant in encoding different tactile cues [16]–[22], e.g., pressure, vibration, shape, texture, the deflection of hair follicles, etc. Moreover, the perception of some elementary cues, such as pressure and flutter/vibration, has been invoked via the intraneural electrostimulation, e.g., slowly adapting (SA) type I and fast adapting (FA) units [23]–[26]. However, device-delivered stimuli do not reflect the full range of naturalistic touch we encounter in everyday life. Indeed, in discriminative touch scenarios, e.g., object manipulation [27] and natural textures [28], that invoke multiple tactile cues, it is likely that single Aβ subtypes provide overlapping and complementary information [29]. Similarly, in human-to-human touch, multiple tactile cues can vary simultaneously [1], [5], [30]. In these types of situations, the analysis of firing patterns becomes much more difficult.

Here, we investigated how the spike firing patterns of Aβ and CT human peripheral afferents encode information about the mechanical inputs produced by human-delivered social touch expressions. Microneurography experiments were conducted as six standardized social touch expressions were delivered. We first characterized afferents’ firing properties, i.e., firing frequency and number of spikes, for comparison to prior studies with well-controlled mechanical contact. Then, machine-learning classifiers were developed to examine the capability of each afferent subtype in differentiating the expressions, for comparison with perceptual studies. Moreover, with these models, we evaluated temporal segments of the full 10 s neural recordings, and their spike-timing sensitivity, to identify the most informative firing patterns for each expression. Overall, the encoding performance of peripheral afferents and their firing characteristics shed light on the information present at the periphery, and which may affect the strategies available to the central nervous system for processing social intent, emotional state or affiliative alignment from physical skin contact.

## Results

### Microneurography paradigm for human-to-human social touch

We developed a novel experimental procedure using microneurography to record single peripheral afferents’ responses to human-delivered social touch expressions. A set of six standardized social touch expressions were delivered by trained experimenters. These standardized expressions were developed based on touch strategies used by people in close relationships, and are recognizable by naïve participants [5], [6] as communicating messages of attention, happiness, gratitude, calming, love, and sadness. Those expressions were specifically performed over receptive fields of identified single afferent units in the bare hand and forearm. Microneurography recordings [31] were obtained from the right radial nerve just above the elbow from 20 healthy participants (Fig. 1A, 1B). All cutaneous afferents were very sensitive to soft brushing and had mechanical (von Frey) thresholds of activation ≤ 1.6 mN. We recorded 39 low-threshold primary afferent units which were classified into six subtypes: fast adapting field (Field, n=5), fast adapting hair follicle (HFA, n=7), fast adapting Pacinian corpuscle (FA-II, n=5), slowly adapting type II (SA-II, n=4), C-tactile (CT, n=6), and muscle spindle (MS, n=12). Per unit, each expression was conducted multiple times, comprising 819 trials in total. All recordings were cropped to keep the first 10 s of data (which was the target duration for the trained experimenters) at a resolution of 1 ms. Examples of collected neural recordings of SA-II and HFA afferents are illustrated in Fig. 1C for all six touch expressions. Despite the consistent delivery of the expressions, distinct firing patterns were observed between these two subtypes. For example, with the sadness and gratitude expressions, SA-II afferents responded throughout contact with a sustained, slowly decaying firing pattern, while HFA afferents only responded to the onset and offset of the holding or when the hand position was adjusted.

**Figure 1.**
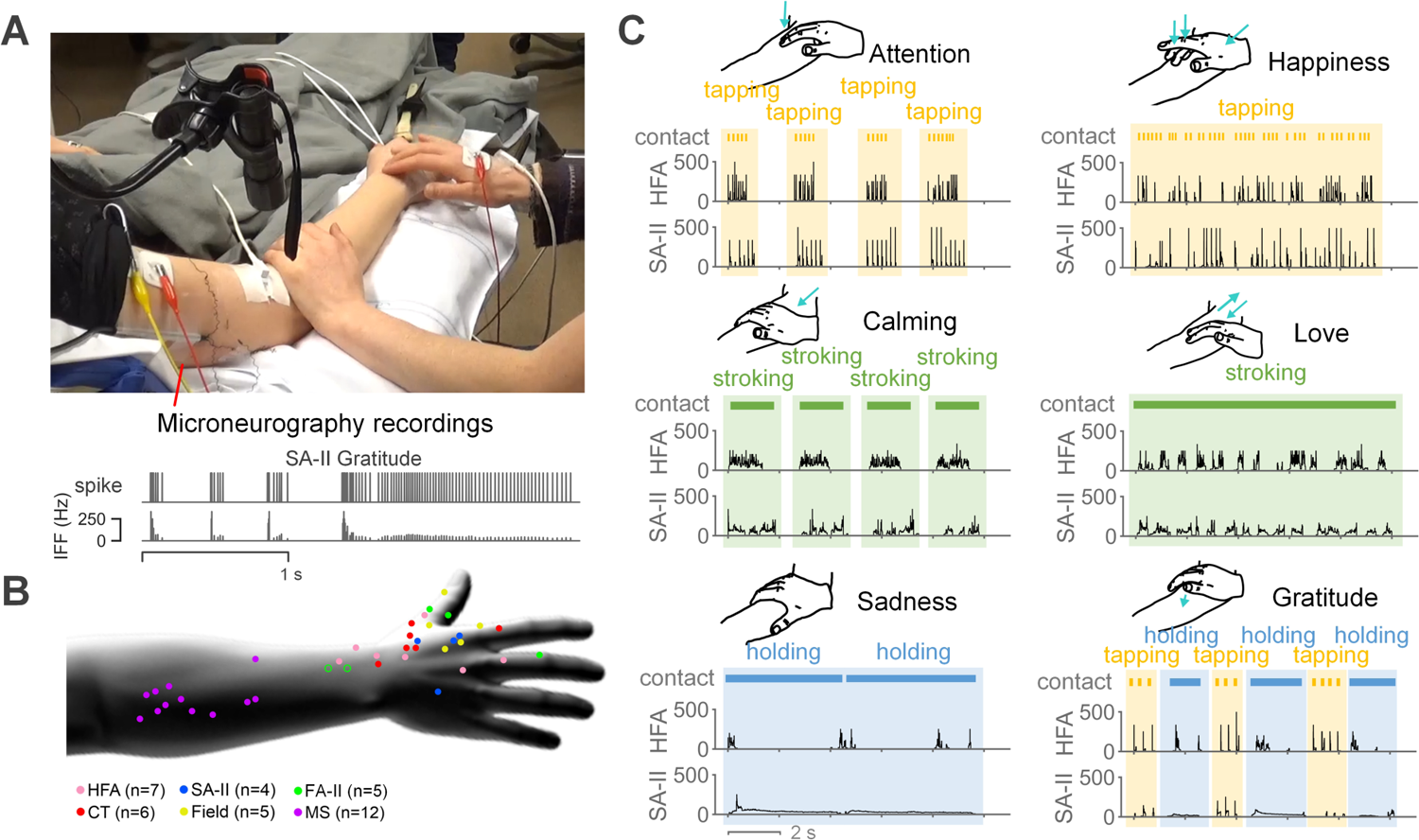
Experimental setup and example microneurography recordings. (A) Standardized touch expressions were delivered over receptive fields of identified afferents by trained experimenters. Microneurography recordings were collected from the upper arm. (B) Multiple units were recorded for each of the six afferent subtypes. For cutaneous afferents, each dot represents the location of an individual receptive field. For two FA-II afferents in the forearm (open circles), the precise location of the receptive field was not documented. For muscle afferents, the dots are shown simply to illustrate where the gestures were delivered. The n-value denotes the number of units per afferent subtype. (C) Six touch expressions were adopted as stimuli. Sketches illustrate the standard contact delivery of those expressions. Component gestures (tapping, stroking, and holding) of the expressions are denoted. Neural response traces illustrate examples of HFA and SA-II responses in instantaneous firing frequency (Hz).

To characterize the firing properties of those afferents in human-delivered touch as compared to controlled device-delivered contact, statistical analyses were performed to quantify and compare mean instantaneous firing frequency (IFF) and the number of spikes across three elementary touch gestures (tapping, stroking, and holding). In particular, attention and happiness expressions were grouped as the tapping gesture, calming and love expressions were grouped as the stroking gesture, and the sadness expression was counted as the holding gesture. Note that the gratitude expression was left out of this analysis since it consisted of both tapping and holding gestures. We combined specific expressions into the three commonly used touch gestures so as to provide a more generalized perspective of afferent’s firing properties, which also align better with the contact interactions examined by controlled stimuli, e.g., indentation, brushing, etc. The results indicate that the observed mean IFF and number of spikes (Fig. 2) share the same ranges with controlled stimuli [10], [16], [19], [32]–[35]. More specifically, the mean IFF of Aβ afferents (around 0-300 Hz) is higher than that for CT and MS afferents (around 0-50 Hz) (Fig. 2A), similar to prior studies using passive touch interactions [10], [32], [35]. For SA-II, HFA, and Field afferents, their mean IFF decreases from tapping to stroking to holding contact, while mean IFF increases for CT afferents (Fig. 2A). Similarly, it has been previously observed that for brush stroke stimuli at velocities above 1 cm/s, the mean IFF for SA-II, HFA, and Field afferents decrease as velocity decreases, while the mean IFF for CT afferents instead increases [10]. As for the number of spikes, rapidly adapting HFA and Field afferents share the same patterns, with stroking contact eliciting significantly more spikes and holding contact eliciting many fewer spikes (Fig. 2B). Note that fewer spikes recorded from tapping contact may be due to the overall shorter contact duration relative to the other two gestures. In comparison, the numbers of spikes for SA-II and CT afferents are also high for slow and static holding contact, which agrees with the firing properties widely reported for these two subtypes [10], [16], [33]. Overall, the spike firing characteristics for the social touch gestures align with those previously identified with controlled stimuli, and help validate the effectiveness of the human touch microneurography paradigm and experiments.

**Figure 2.**
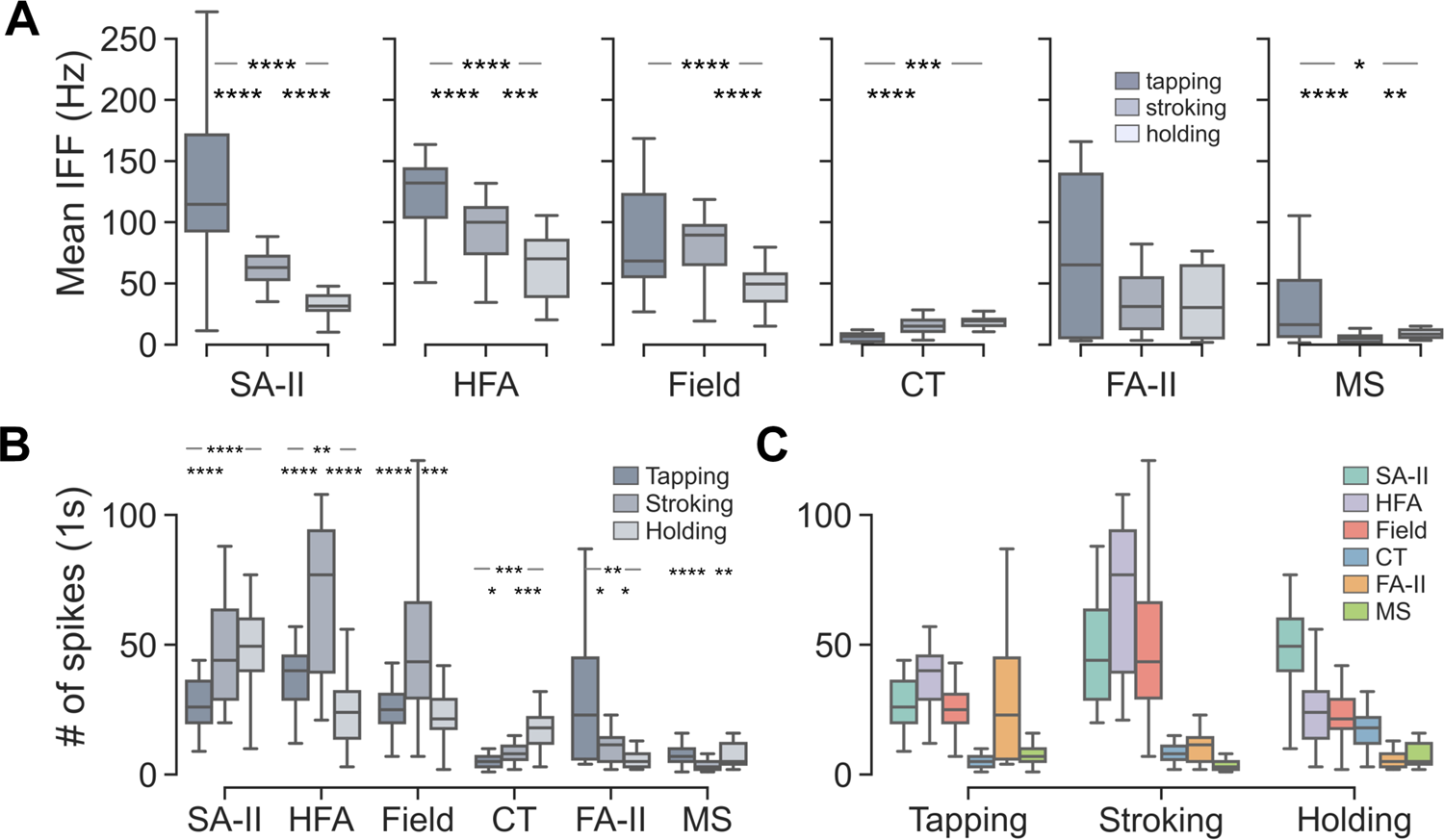
Comparison of mean instantaneous firing frequency (IFF) and the number of spikes across afferent subtypes and gestures. (A) Distributions of mean IFF across gestures per afferent subtype. (B) Distributions of the number of spikes across gestures per afferent subtype. The number of spikes per trial was calculated from a 1 s duration with the largest number of spikes. (C) Distributions of the number of spikes across afferent subtypes per gesture. Significance test results are not shown in this panel and can be found in Fig. S1. \**p* < 0.05, \*\**p* < 0.01, \*\*\**p* < 0.001, \*\*\*\**p* < 0.0001 were derived by Mann–Whitney U tests with Benjamini-Hochberg post-hoc correction.

More interestingly, these peripheral afferents well differentiate the three touch gestures based upon only the two firing features evaluated (Fig. 2A, 2B). Indeed, despite contact variations tuned by social meaning, the contact patterns of those gestures are distinct enough to elicit reliable and significantly different firing patterns (Fig. S1B). That said, discrepancies are observed among the firing properties across afferent subtypes. For example, CT afferents exhibit a different trend of mean IFF across gestures compared with Aβ afferents; the number of spikes of SAII afferents follows a different trend than that of HFA and Field afferents. Moreover, FA-II and MS subtypes provide relatively less information in encoding gestures, which may relate to the extremely high sensitivity of Pacinian corpuscles [19] and the proprioceptive functionality of muscle spindles [35]. Moreover, while all Aβ afferents responded very well to tapping contact (Fig. 2C), SA-II responded with significantly more spikes for holding and FA-II exhibits significantly fewer spikes for stroking. These distinct properties suggest the potential for complementary functional roles of those afferents when viewed as a population at higher levels of the nervous system.

### Single units of SA-II and HFA afferents effectively encode social touch expressions

In order to evaluate how well different classes of primary afferents are able to discriminate the six expressions, we developed a time-series classifier using a one-dimensional convolutional neural network (1D-CNN) to predict delivered social touch expressions from the neural spike trains. The model was trained and tested for each afferent subtype separately with full 10 s binary spike trains fed as input. SA-II and HFA achieve the highest prediction accuracies around 70-80% (Fig. 3A). Note that the results may slightly vary due to the random train-test splitting and stochasticity of CNN model. Such accuracies are very close or even slightly higher than human recognition accuracy with the same six standard touch expressions [5]. In comparison, Field afferents afford relatively lower accuracy around 56%, while the accuracy of CT, PC, and MS afferents are not far from the chance level of 16.7%. In summary, this analysis indicates that SA-II and HFA subtypes convey the richest information among the tested six afferent subtypes in encoding the six social touch expressions.

**Figure 3.**
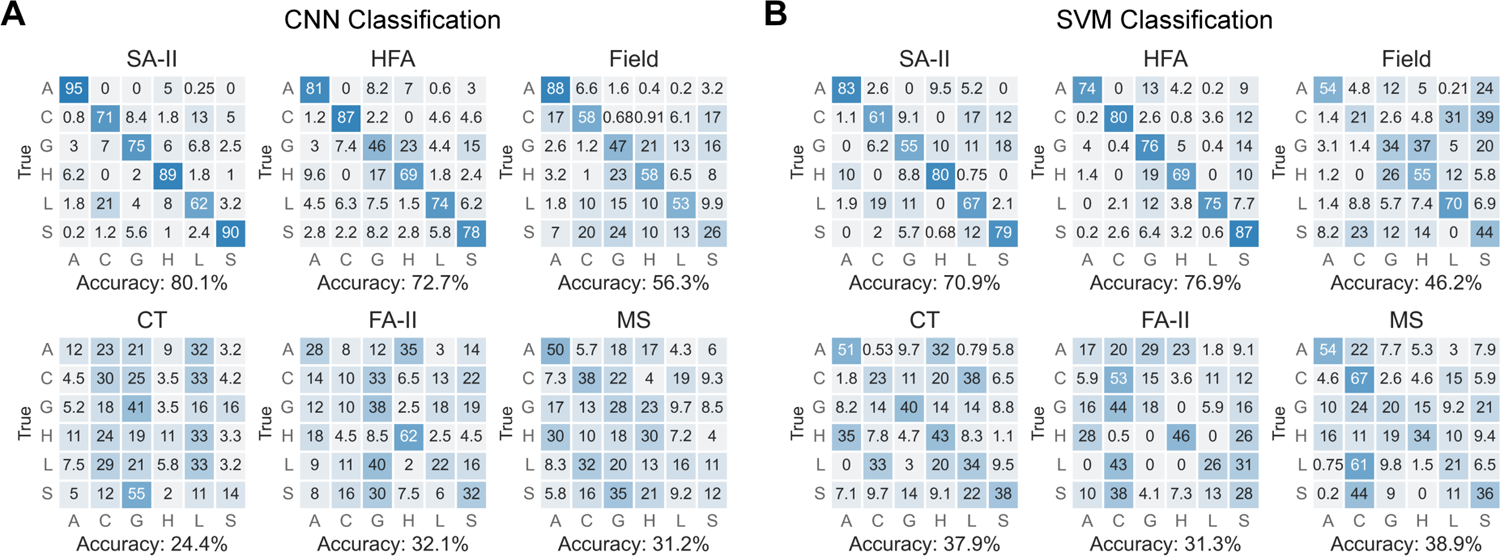
Expression classification using CNN and SVM models. (A) Classification results per afferent subtype using the CNN classifier with 10 s spike trains as input. (B) Classification results per afferent subtype using the SVM classifier with five features extracted from 10 s recordings as input. Across both models, the classification accuracy is markedly higher for SA-II and HFA subtypes, at levels observed in human perceptual experiments [5].

To better examine the encoding capability of those afferents, a simpler model in the form of a linear support vector machine (SVM) was employed for classification again to decouple the computational power of the more complex CNN model. Instead of an input of time-series neural recordings, an input of five features extracted from 10 s recordings was used, which includes mean IFF, number of spikes, peak IFF, IFF variation, and number of bursts (details in Methods). As shown in Fig. 3B, similar to the results with the CNN model, high classification accuracies of around 70-80% were observed for SA-II and HFA subtypes. Also, in alignment with results obtained with the CNN model, the Field, CT, FA-II, and MS afferents exhibit lower accuracy. The consistency in classification performance between the two models implies that the identified encoding power of those afferent subtypes is associated with the information carried by their firing patterns instead of being a function of the model itself. Among the six subtypes, SA-II and HFA afferents are capable of encoding social touch expressions in an accurate and reliable way.

### Most informative firing patterns for encoding social touch expressions

In order to identify the most informative segments of firing patterns that lead to high differentiation accuracies, we further conducted CNN classification on segments of neural recordings per afferent subtype. A sliding window method incorporating window position (Fig. 4A) and window length (Fig. 4C) was applied to segment chunks from a given train of neural recording for comparison (details in Methods). Indeed, the length of neural responses has been well characterized in encoding discriminative touch with controlled stimuli [16]–[22]. In contrast, here we are interested in exploring and comparing the window length of neural recordings in encoding human social touch, which affords a higher level of spatiotemporal complexity. 64 different window lengths were selected ranging from 0.1 to 10 s. For each window length, five segments at different positions were derived according to five metrics using sliding window method with the step of 1 ms, which includes the first segment, the segment with the largest number of spikes, the segment with the highest mean IFF, the segment with the highest IFF variation, and the segment with the highest IFF entropy. Therefore, 320 different segment options were obtained in total and were compared using the CNN model in terms of classification accuracy.

**Figure 4.**
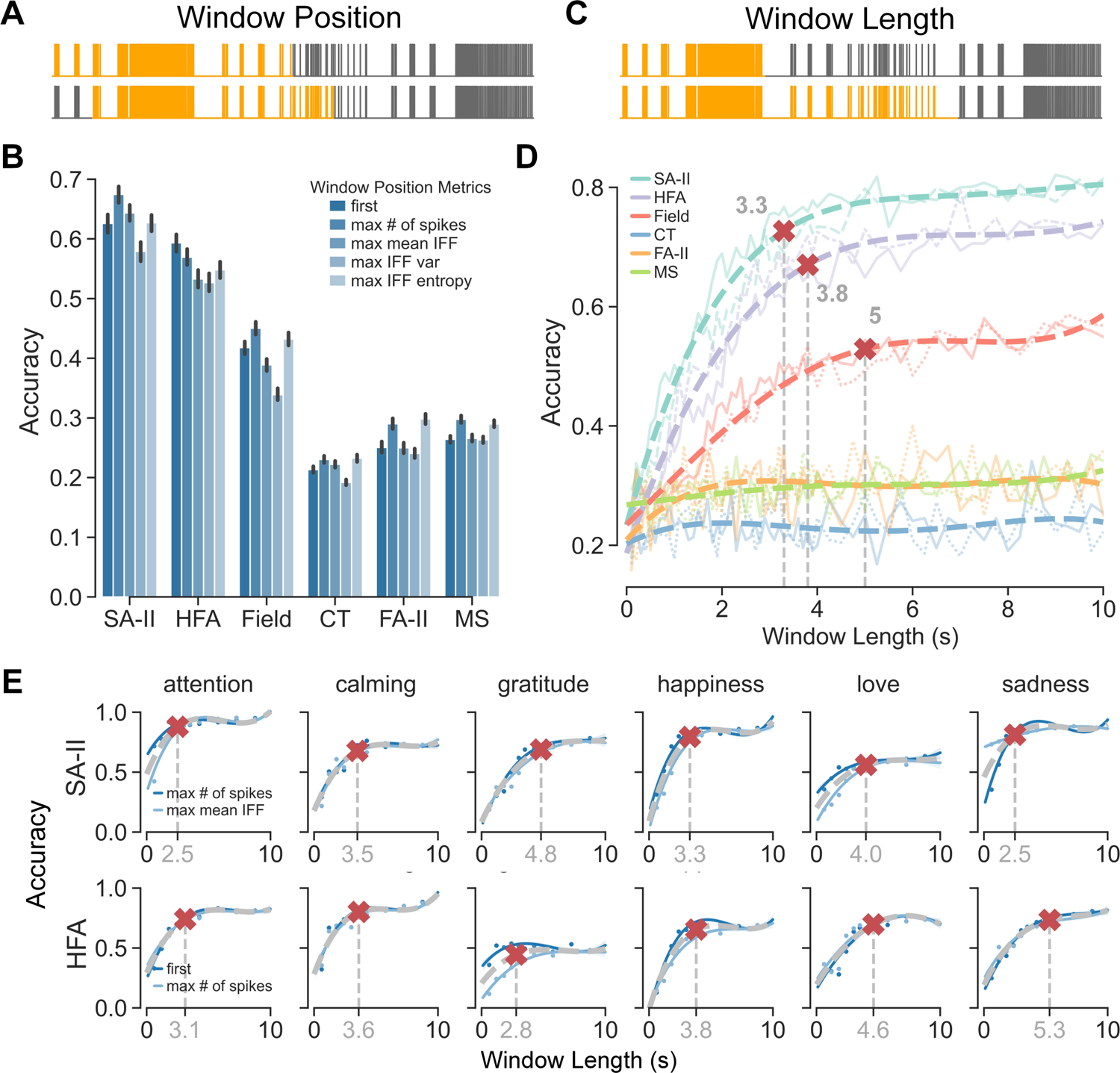
Comparison of CNN classification accuracies when using segments of the full 10 s neural recordings derived from different window positions and window lengths. (A) An example of two window position options with the same window length. Gray traces are 10 s spike trains from the same trial, where highlighted spikes illustrate two different segments. (B) Classification accuracies across window position metrics averaged over all window lengths for each afferent subtype. Significance test results are not shown in this panel and can be found in Fig. S2. (C) An example of two window length options with the same window position. Gray traces are 10 s spike trains from the same trial, where highlighted parts illustrate two different segments. (D) Classification accuracies with window length per afferent subtype. Accuracy curves were fitted using data from their best two window positions and fourth-order polynomial regression, shown as dotted curves. Red cross markers denote 90% saturation window lengths. Two lighter curves represent data from the two best two window positions. (E) Classification accuracies along with window length per afferent subtype per expression. Averaged accuracies from their best two window positions are shown as grey dotted curves and blue curves represent each of the best positions. Red cross markers denote 90% saturation window lengths.

The five window position metrics were first compared per afferent subtype with all window lengths combined. For most pairs of metrics, significant differences were identified in average classification accuracy (Fig. 4B, details in Fig. S2). However, the differences in accuracy are only around 3.3% between the top two metrics for SA-II afferents (the highest number of spikes and the highest mean IFF) and around 2.7% for HFA afferents (first and the highest number of spikes). Moreover, five accuracy curves along with the window lengths corresponding to the five metrics also well overlap, especially for SA-II and HFA subtypes (Fig. S2). The overlapping curves indicate the major impact of window length on classification performance with window position leading to a lesser difference.

Among the five window position metrics, the two top performing metrics were adopted for use in examining the influence of window length. In general, the results indicate that classification accuracies for SA-II, HFA, and Field afferents saturate when window length increases (Fig. 4D). In contrast, accuracies for the other afferent subtypes begin and remain consistently low. For SA-II, HFA, and Field subtypes, we identified the saturation window lengths from their accuracy curves based on 90% of their highest accuracies using fourth-order polynomial regression, which are 3.3 s, 3.8 s, and 5 s respectively. It implies that instead of the full 10 s, an average duration of 3-4 s of the neural responses of SA-II and HFA afferents provide sufficient information to differentiate the expressions. Accuracy curves and saturation window lengths were further identified for SA-II and HFA subtypes and all expressions (Fig. 4E). Variation between 2.5 to 5.3 s was observed among the identified saturation window lengths, which is still a comparably limited range. It again supports that much less duration from the full 10 s is required for SA-II and HFA afferents to encode the social expressions.

According to best window positions and saturation window lengths of SA-II and HFA afferents, the most informative segments of their firing patterns are identified for each expression (Fig. 5A). Note that for expressions with multiple rounds of contact, e.g., attention and calming, prolonged non-contact/non-response gaps are included, which ties to the nature of the rhythm of the toucher’s contact delivery. More interestingly, it illustrates that the variation of saturation window length across expressions could be related to both contact rhythms of touch expressions and firing properties of afferent subtypes. For example, the unique repetitive tapping pattern of attention expression might explain why it requires relatively less data than other expressions. Sadness exhibits the largest difference in saturation window length between SA-II and HFA (Fig. 4E). One explanation is the sustained low-frequency firing pattern of SA-II afferents under holding contact is easy to differentiate even within a shorter time. In comparison, the firing pattern of HFA to holding is similar to that of tapping contact such that more data including the non-response gap are needed to capture the unique dynamic of prolonged holding of the sadness expression (Fig. 5A).

**Figure 5.**
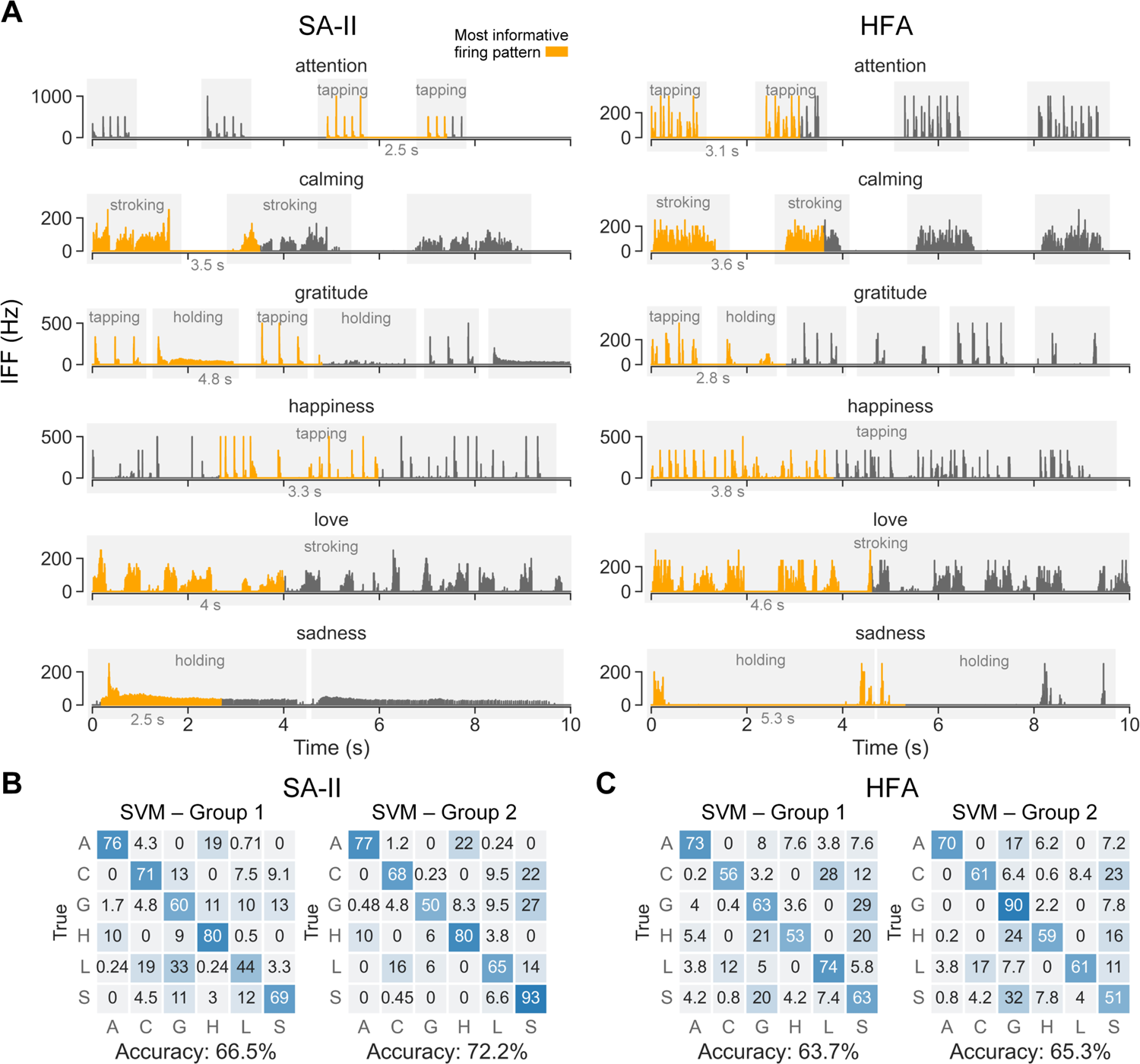
Examples of identified most informative firing patterns and SVM classification based on the identified segments. (A) IFF traces highlighted in orange are the most informative segments determined by the best window position (SAII: max # of spikes, HFA: first) (Fig. 4B) and the saturation window length per afferent-expression combination (annotated under the highlighted segments) (Fig. 4E). Grey shades represent gestures of tapping, stroking, and holding. (B, C) Classification results using the SVM model. Group 1 and group 2 refer to two groups of neural recording segments derived by saturation window lengths per afferent subtype (Fig. 4D) and saturation window lengths per afferent-expression combination (Fig. 4E), respectively. (B) results for SA-II subtype, (C) results for HFA subtype.

The identified segments of SA-II and HFA recordings were then classified using the SVM model to compare with the performance of using full 10 s recordings. Two groups of segments derived from saturation window lengths per afferent (Fig. 4D) (group 1) and per afferent-emotion combination (Fig. 4E) (group 2) were classified (Fig. 5B, 5C). The best window position metric (Fig. 4B) was applied to both groups. As those window lengths and positions were identified using the CNN model, classification with just those segments provided similar accuracy results as those with the full 10 s recordings, as expected (Fig. S3). Meanwhile, classification accuracies using the SVM model are also similar to the full 10 s recording results (Fig. 5B, 5C), where slightly higher accuracies were obtained for group 2. Such findings validate the richness of information contained within identified segments of SA-II and HFA afferents’ firing patterns.

### Spike-timing sensitivity in human social touch

As an additional way to examine the temporally relevant features of the spike train, we examined the spike-timing sensitivity of the SA-II and HFA subtypes in classifying the social expressions. To achieve this, random noise was added to the spike times across the 10 s neural recordings used by the CNN classifier (Fig. 6A). Noise following a Gaussian distribution was employed with mean equals to zero and standard deviation (SD) ranges from 0 to 100 ms with a step of 5 ms. The CNN model was trained per subtype with noise-less neural recordings and was tested using recordings with noise added. Among the six afferent subtypes, the SA-II and HFA subtypes were most sensitive to spike-timing noise as their classification accuracies drop drastically when noise increases (Fig. 6B). However, they also exhibited tolerance to lower noise at around 10 ms SD. This tolerance could relate to the variability of the human-delivered touches, the variability of firing patterns across different units, and/or the aggregated information of expression as the prediction target. In contrast, spike-timing noise caused little impact on classification performance of the other afferent subtypes. Moreover, for SA-II afferents, the attention, happiness, sadness and gratitude expressions were sensitive to noise (Fig. 6C), which were all delivered by tapping or holding contact. HFA afferents were sensitive to noise with the attention and happiness expressions, which were only delivered by tapping. This difference aligns with the firing property of SA-II afferents that they respond to holding with a distinct, sustained firing pattern. Except for them, all other expressions and afferent combinations were robust to noise across tested noise levels.

**Figure 6.**
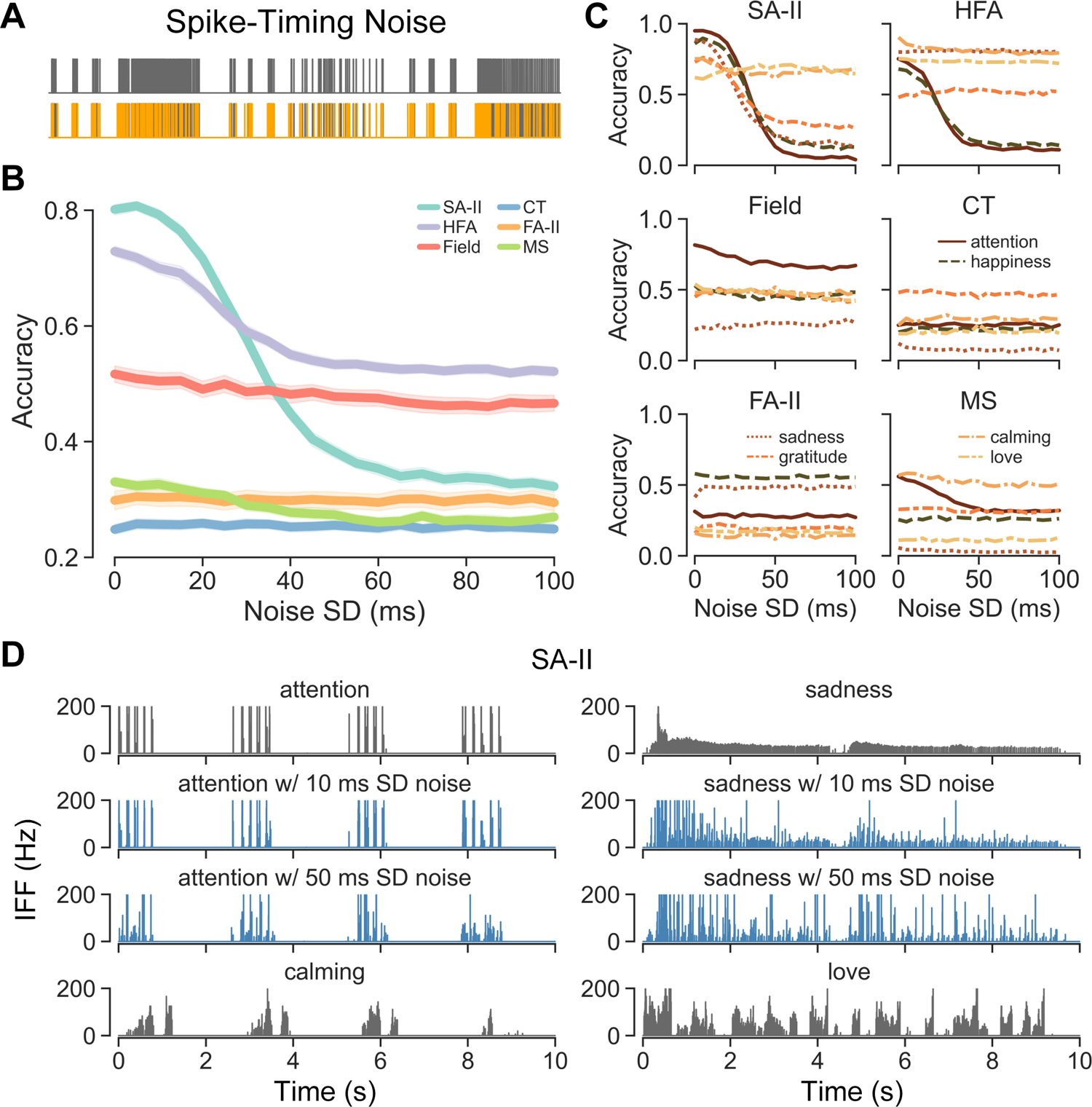
Spike-timing sensitivities across afferent subtypes in human social touch. (A) Spike trains from one trial of gratitude expression with (lower) and without (upper) spike-timing noise added. (B) CNN classification accuracies relative to the standard deviation of added noise. (C) CNN classification accuracies of expressions relative to the standard deviation of added noise. (D) Two confusion cases with SA-II afferent recordings. The IFF traces in grey are the original neural recordings and IFF traces in blue are the neural recordings with 10 ms and 50 ms standard deviation noise (SD) respectively. Attention expressions were misclassified as calming and sadness expressions were misclassified as love when 50 ms SD noise was added.

To investigate the potential cause of such high spike-timing sensitivity of certain afferent-expression combinations, neural recordings with and without noise were compared in those confusion cases. According to the confusion matrix of SA-II afferents with 50 ms SD noise, attention was misclassified as calming and sadness was misclassified as love (Fig. S4). While 10 ms SD spike-timing noise did make a huge difference on firing patterns, we found that noise as high as 50 ms SD could flatten out isolated spikes elicited by repetitive taps within one round of tapping (Fig. 6D). It thus changed the envelope of the firing pattern to be a continuous chunk of firing with variable frequencies, which is similar to the firing pattern of stroking contact. As for the holding contact of sadness, spike-timing noise could convert its sustained slowly decaying firing pattern into a spiky and irregular shape, similar to the firing pattern of stroking (Fig. 6D). Here, attention and calming were mainly confused with the stroking of calming and love respectively, which could relate to their shared touch rhythm of having prolonged non-contact gaps, or not. Based on the above observations, we hypothesize that the spike-timing sensitivity of those afferent subtypes could be strongly tied to the extent of changes in the envelope shape of their firing patterns caused by noise. This envelope information might be better at capturing the contact pattern at a macro level, such as gestures, to encode touch expressions. In this scenario, millisecond-precision of single spike times might not be as informative due to robustness of the touch expressions and their social meanings.

## Discussion

We might lightly tap someone to get their attention or stroke a partner’s arm to offer a sense of calm. How these expressions are perceived is influenced by how peripheral afferents encode mechanical contact. Up to now, the properties of touch afferents have been exclusively characterized with precisely-controlled stimuli (e.g., vibration, force-calibrated indentations or tangential loading of the skin). In this work we explore neural responses from multiple types of afferents to human-delivered social touch. Overall, the findings indicate that single units of Aβ afferents, especially SA-II and HFA subtypes, can readily differentiate social touch expressions, at perceptually-observed levels. Moreover, the analysis of spike firing patterns, using conventional statistical analyses and classification analyses using convolutional neural networks and support vector machines, indicates that temporal firing envelopes of about 3-4 s and spike-timing precision of 10 ms afford sufficient discriminative information.

### Microneurography paradigm for human-to-human touch

Distinct from traditional experiments that control the mechanical stimulus and vary a single feature at a time, we record from single peripheral afferents in a human-to-human touch paradigm, where multiple stimulus features, e.g., normal displacement, contact area, lateral velocity, vary simultaneously [1], [5], [36], [37]. Such naturalistic social touch interactions are closely tied to our human well-being and development, and may extend into everyday tasks such as feeding, grooming [38], and caregiving [39]. However, these types of interactions are technically difficult to replicate with actuated devices. Indeed, precisely controlled stimuli, such as rigid bodies indented in one dimension of depth or force [17]–[19], are more commonly employed in characterizing the firing properties of peripheral afferents. Recent efforts have begun to move toward more naturalistic contact interactions using brushing, puffs of air, and pinch, etc., [40], [41]. Natural textures have also been applied in recording monkey Aβ afferents [28]. However, each of these efforts still controls and varies a single stimulus feature at once, which is different from natural contact with co-varying features. In this study, we move a step further into human-delivered touch, where the richness of contact dynamics could reveal classes of primary afferents that encode the combination of multiple features. In our tasks, such information could be relevant to social messages conveyed in touch expressions. More specifically, six standardized social touch expressions were delivered by trained experimenters. This affords reliable contact interactions [5] and retains the subtleness of human-delivered touch at the same time. Meanwhile, expressions were designed with specific touch gestures, which can be compared with similar mechanical stimulus contact, e.g., human-delivered stroking versus brush-delivered stroking, human-delivered tapping versus vibrating actuator indentation. Indeed, the firing properties we observed in human touch (Fig. 2) share similar ranges and trends with those for controlled stimuli. It also demonstrates that similar states of skin contact and deformation could elicit similar responses across human touch and stimulus contact [42].

### Social touch encoding across afferent subtypes

Two afferent subtypes, SA-II and HFA, stand out in their ability to differentiate the six social touch expressions. In particular, both CNN classification using time-series neural recordings and SVM classification using five firing features show that those two subtypes outperform other subtypes (Fig. 3) and provide high differentiation accuracies similar to human perception [5]. Moreover, such accuracy is consistent in using either the full 10 s time course of the neural responses or the most informative firing patterns therein (Fig. 5B, 5C).

The SA-II and HFA afferent subtypes, due to particular physiological mechanisms, may be geared more to the inherent contact characteristics of the six touches. For example, one prominent commonality between these two subtypes is their large, but not too diffuse, receptive fields [43]–[45], which may help in consistently capturing the range of contact dynamics given the size of touchers’ fingers and hands and their lateral movement patterns. In particular, rapidly adapting units exhibit large receptive fields in the hairy skin of the human forearm [44], which are 78-113 mm^2^ as opposed to 24 mm^2^ on the dorsum of the hand [46] and 13 mm^2^ in glabrous skin [43]. SA-II afferents exhibit receptive fields around 28 mm^2^ in the dorsal hand [46] and are found to increase in size considerably with indentation force, as compared to SA-I units [44]. FA-II afferents that innervate Pacinian corpuscles may exhibit too large of receptive fields, which are almost too diffuse to map due their extreme sensitivity [19]. Moreover, FA-II afferents are relatively scarce in hairy as opposed to glabrous skin [46]. Therefore, compared with other subtypes, the relative size of the receptive fields of HFA and SA-II afferents in hairy skin could contribute to their social expression encoding.

Furthermore, SA-II and HFA afferent subtypes are believed to be sensitive to a wide range of contact, including hair deflection [47], skin stretch, and shearing forces [16], [48], which are contact characteristics that human touch gestures tend to evoke. For example, both SA-II and HFA respond to tapping (vertical contact) and stroking (sheering contact) with distinct mean IFFs (Fig. 2A) and can easily differentiate those two gestures (Fig. S1). In contrast, Field and FA-II afferents respond to these two directions of contact with non-differentiable firing frequencies (Fig. 2A). Moreover, both SA-II and HFA afferents precisely followed tapping contact with high IFF responses (Fig. S5), outperforming the other subtypes. Interestingly here, SA-II afferents are typically thought to mainly encode static/slow movements and skin stretch [16], [48], but also responded very well to fast vertical contact delivered by human tapping. As for holding contact, as expected, SA-II afferents respond with sustained low-frequency firing patterns, which distinguish holding from other fast movements. HFA afferents did not respond to the sustained contact, but precisely captured the on-set and off-set of the hold gesture. Although this pattern of spike firing is similar to that of tapping, the unique prolonged touch rhythm of holding provides distinct temporal information (Fig. 1C). Meanwhile, the mean IFF in the case of the holding gesture is significantly lower than for other gestures, which may relate to slow movements and gentleness deployed in holding contact. Overall, the capability of SA-II and HFA subtypes to differentiate the social touch expressions suggests that their neural responses well correspond to the range of stimulus input and mechanical skin deformation inherent in human-to-human touch interactions.

Focusing on the context of social touch, the afferent subtypes exhibited distinct sensitivities in encoding the two layers of information, i.e., gestures (lower level) and expressions (higher level). Based on the same five firing properties, all six subtypes could accurately differentiate the three gestures (Fig. S1), whereas CT, FA-II, and MS afferents fail to separate the expressions (Fig. 2B). It suggests that the distinct contact patterns of elementary touch gestures, e.g., tapping, stroking, and holding, can be captured to a certain extent by all afferent subtypes. Indeed, and in comparison, expressions delivered by selective use of the same gesture can be fine-tuned to convey specific social meanings by participants making subtle changes in skin-to-skin contact, e.g., velocity, indentation depth, contact area [36]. Such nuances may be less easy to capture for certain afferents. For example, attention and happiness, delivered by tapping, and calming and love, delivered by stroking, were frequently confused by CT, FA-II, and MS subtypes. From another perspective, it is also reasonable that those pairs of expressions might be confused as they share relatively similar contact dynamics, and are also likely to be misidentified by human receivers [5]. On the other hand, SA-II and HFA subtypes are very sensitive to slight contact changes, as is observable in the lack of a discrepancy between their gesture and expression classification accuracies (Fig. S1, 2B).

The relatively lower coding capability of CT, FA-II, and MS afferents might be linked to their functional roles in signaling specific contact modalities less reflective of the differences in the six social touch expressions. Among these subtypes, CT afferents are traditionally thought to be associated with affective touch, more specifically pleasantness as elicited through stroking [8], [49]. It has been suggested that CT afferents respond preferentially to certain contact patterns and velocity ranges related to the hedonic processing, in parallel with Aβ afferents serving as discriminator for different contact stimuli [50]. In alignment with such roles, we indeed found that CT afferents can successfully identify stroking contact among the six examined touch expressions (Fig. S1), yet could not further differentiate contact differences between the expressions of love and calming (Fig. 2B). Those two expressions are presented as similar gentle stroking with different stroking routes and contact rhythms, i.e., love: continuous back-forth stroking, calming: four separate one-direction stroking. The lack of discrimination of those contact differences aligns with reports of low spatial and temporal sensitivity of single CT units [51], although a population of CT units may perhaps better inform such affective sensation [52]. Meanwhile, holding contact tends to be misclassified as stroking, which is reasonable since slight hand movements always exist among human delivered touch. Surprisingly, CT afferents also respond very well to fast vertical tapping contact (Fig. S5). Although CT units have been reported to respond well to von Frey indentation [53], human tapping affords much higher levels of force in a faster and repetitious manner. However, within the same tapping gesture, more detailed contact differences between expressions of attention and happiness were not captured in responses of CT afferents. For the other two subtypes, FA-II afferents are respond to high-frequency vibration in discriminative touch, such as contact delivered to a site remote to the center of the afferent’s receptive field center [46]. However, they filter low frequency stimuli [19] that carry most of the information adhering to social touch. MS afferents respond to muscle extension and flexion associated with our proprioceptive mechanism [54], which means they are less likely to be activated when being passively touched by human touchers.

### Temporal envelope of firing pattern as potential strategy of social touch perception

By leveraging machine learning classification models, we identified the most informative firing patterns of SA-II and HFA afferents in encoding touch expressions. Those firing patterns and their corresponding contact patterns suggest strategies tied to similar levels of perceptual discrimination. More specifically, instead of the full 10 s time course of contact, we found that an average of 3-4 s provides enough information for single units to differentiate the six expressions (Fig. 4D). Also, as window position did not have a critical impact, it suggests that afferents respond in a consistently informative way throughout the course of contact, where the accumulation of a sufficient amount of information would be the key for social touch processing. Indeed, this time duration of 3-4 s aligns with the cortical response time of brush-delivered affective touch [55], facial EMG response time in natural social touch that reflects emotional processing [5], and the acceptable response time of humanoid robots being touched by a human [56]. However, this time duration is significantly longer than that reported in encoding precisely-controlled single stimulus features. Based on a population simulation of peripheral tactile afferents, tens of milliseconds were found to be sufficient in encoding stimulus directions [57]. Similarly, with ensembles of recorded single afferents, tens of milliseconds neural responses were also suggested to be effective in encoding controlled force, torque, force direction, and shape on finger pads [58], [59]. This difference in neural response duration for touch encoding may highlight the complexity of human social touch, where social meaning may be integrated from varying combinations of spatial and temporal contact interactions. For example, attention is generally expressed as several rounds of repeated fast tapping versus happiness is expressed as continuous tapping with fingers dancing back and forth on the receivers’ forearm. Tapping contact from the two expressions might be signaled by peripheral afferents in a comparable way within tens of millisecond duration. Nevertheless, they could be differentiated from firing patterns reflecting touch gestures and contact rhythms in the time scale of multiple seconds (Fig. 5A). On the other end of the spectrum, controlled stimulus features typically carry less information of constant states, e.g., a certain direction of indenting stimulus, and thus could be identified with much shorter neural responses especially using a population model [57]. As single units were tested individually in our case, we expect that population responses of single or multiple afferent subtypes might encode social touch expressions in shorter durations.

Furthermore, peripheral neural responses allow for the precise timing of the spikes to be shifted by about 10 ms with little effect on the classification of the expression, although greater shifts can change the firing patterns so that an expression is confused with another. It appears that the spike timing precision needed in encoding human social touch is relatively lower than encoding traditional stimulus features. For instance, when classifying well-controlled scanned textures and vibratory stimuli, the optimal spike timing precision is around 1-10 ms [28], [60]. Yet consistent with these reports, we also find that slowly adapting afferents are less sensitive to spike timing than fast adapting afferents. In the above studies [28], [60], the distance to transform one spike train to another [61] was used to classify replicated stimuli. As some variation can exist in the exact human contact delivery, we directly added artificial jitter to spike times [62]–[64] and tested them with the high-resolution time-series CNN model. It was believed that spike-timing jitter would blur the transmitted information of the stimulus [62], [63], [65]. For encoding controlled audio amplitudes, milliseconds or even sub-milliseconds of added artificial jitter can significantly decrease the accuracy of transmitted information [64]. Therefore, in human social touch interactions, the relatively higher tolerance to spike-timing jitter suggests that the aggregated temporal pattern might play a role in capturing the delivered expression information. We found that by adding spike timing jitter, the envelope of the firing pattern can be drastically changed. Through qualitative observation, such coarse-grained temporal patterns may be closely related to macro level information, e.g., touch gestures (Fig. 6D), instead of cell level dynamics of signal transmission. It also aligns with the finding that the SVM model using aggregated firing properties provided comparable classification performance as the time-series CNN model (Fig. 2A, 2B, 5B, 5C). Their similar performance implies that detailed spike-to-spike temporal coding may not contribute to the core information in complex social touch scenarios. On the other hand, rate coding of statistical features could capture the temporal pattern to a certain extent but might not capture the whole dynamics. Here we hypothesize that the temporal envelope of the firing pattern, which falls between the precise temporal coding and the rate coding, could be a valuable metric in representing social touch expressions, where the window length would also have a large impact.

### Limitations and future works

The slowly-adapting type I (SA-I) afferent is another Aβ subtype that is likely to play a significant role in encoding social touch stimuli. In general, SA-I afferents contribute to our abilities in fine touch discrimination, as demonstrated in experiments with precisely-controlled stimuli [45], [66]. In our study herein, the population of SA-I afferents (n=2) was not large enough to include. Our speculation is that SA-I afferents might behave akin to SA-II afferents, due to similar adaptation characteristics. Additionally, SA-I afferents exhibit a very large dynamic range of sensitivity, as compared to SA-II afferents, combined with very low absolute thresholds [67]. On one hand, such sensitivity should benefit discriminability, in general. On the other hand, in discriminating social touches, if SA-I afferents are too sensitive, this may be too variable a response that buries the core contact information carried by the temporal envelope of the firing pattern. In this way, SA-II afferents might offer advantages because they have relatively dampened responses to dynamic stimuli, compared to SA-I afferents. Indeed, FA-II subtype that exhibits extremely high temporal sensitivity provides low differentiation accuracy. Work with high threshold mechanoreceptors has also shown that less sensitive subtypes can be better encoders of noxious forces than those relatively more sensitive [24]. However, further follow up work is required to understand the response characteristics of the SA-I subtype to social touches.

Additionally, at the single-unit level, it is possible that SA-II and HFA afferents may struggle to distinguish different sets of touch expressions than those we used, and other subunit types may excel. We designed the expressions to cover a wide range of naturalistic interpersonal touch interactions that are understandable by human receivers. Meanwhile, the designed expressions were connected to specific social meanings so that the underlying emotional contexts could be moderated. In particular, the perception of pleasantness (valence), emotional arousal, and dominance [68], [69] were not fully explored in this study. Part of the reason was to avoid the high task load of participants if psychophysical and microneurography experiments were conducted together. Based on the dataset of emotional ratings for English words [69], we found that happiness and attention are particularly high arousal and were both delivered by fast tapping contact. We might assume that neural responses to fast contact velocities are related to high arousal percepts. However, other contact characteristics, e.g., force, indentation, contact area might also contribute [36]. Therefore, precise contact quantification needs to be introduced to uncover further details of how emotional contexts of physical touch delivery are encoded by peripheral afferents [37].

Moreover, as we hypothesize that the temporal envelope of firing pattern could suggest the potential strategy of how humans perceive social meanings from touch, it needs to be further assessed in a systematic way. On top of that, the population encoding mechanism is also not fully understood for social touch interactions. More specifically, while single units appear to hold discriminative capacity, afferent subtypes are likely to interplay in a cohesive way in generating population responses [29], [70], from which our perception and discrimination are gleaned. Our findings regarding single unit responses provide the foundation for such future explorations, where empirical or mathematical studies of higher-order nervous structures would benefit our understanding of population processing of social touch communication.

## Methods

### Participants—touch receivers

All participants were recruited through local advertisement and a mailing list. Using the microneurography technique for single-afferent axonal recordings, responses to social-touch expressions were recorded from 41 low-threshold primary afferent fibers belonging to the right radial nerve in 20 healthy participants (23-35 years old – all except 1 who was 50 years’ old; 13 males, 5 females). All participants provided informed consent in writing before the start of the experiment. The study was approved by the ethics committees of Linköping University (Dnr 2017/485-31) and complied with the revised Declaration of Helsinki. The participants were seated in a comfortable chair and pillows were provided to ensure minimal discomfort.

### Standardized touch expressions

Based on observations of the common physical features of touch communication behavior between people in a close relationship [5], we developed a set of standardized touch expressions to communicate “attention,” “happiness,” “calming,” “love,” “gratitude” and “sadness.” More specifically, the touch expression of “attention” comprised 4 bursts of 4-5 repetitive taps with the index finger, each burst lasting approximately 1.5 s, with approximately 1 s between. “Happiness” consisted of continuous random playful tapping, using multiple fingers, and moving up and down the arm. “Gratitude” consisted of patting (3-4 pats with multiple fingers, lasting approximately 2 s) alternated with holding (long grasp with the whole hand, lasting approximately 2 s). “Calming” involved 4 repeated strokes down the arm with the whole hand, each lasting approximately 2 s, with approximately 0.5 s between. “Love” involved a continuous back-and-forth stroking with the fingertips up and down the arm. Finally, sadness consisted of a sustained hold on the arm with light squeezing.

These standardized expressions were applied by trained experimenters to the physiological receptive field of single neurons during microneurography recordings. The experimenter received spoken cues via headphones, first the cue-word, then a countdown (3, 2, 1, go). They were instructed to perform the touch starting from the “go”-signal until they heard a stop signal (3, 2, 1, stop), creating a continuous time window of touch for 10 s. The experimenter was first familiarized with the afferent’s receptive field and was required to touch an area of skin including but not limited to the receptive field. They were also required not to perform any vigorous movements to avoid dislodging the recording electrode. Where a single-unit recording was stable enough, data for multiple trial-sets were obtained.

### Microneurography

Neural recordings were performed with equipment purpose-built for human microneurography studies from ADInstruments (Oxford, UK; setup 1) or the Physiology Section, Department of Integrative Medical Biology, Umeå University (setup 2). The course of the radial nerve just above the elbow was visualized using ultrasound (LOGIQ e, GE Healthcare, Chicago, IL, USA). A high-impedance tungsten recording electrode was inserted percutaneously and with ultrasound guidance it was inserted into the nerve. Where needed, weak electrical stimuli through that electrode were delivered to localize the nerve (0.02-1 mA, 0.2 ms, 1 Hz; FHC, Inc. Bowdoin, ME, USA). The electrode was insulated, except for the ∼5 µm bare tip, with a typical length of 40 mm and shaft diameter of 0.2 mm. In addition to the recording electrode, an indifferent (uninsulated) electrode was inserted subcutaneously, approximately 5 cm away from the nerve. Once the electrode tip was intra-fascicular, minute movements were made to the recording electrode, manually or with a pair of forceps until a single afferent signal was isolated. Each low-threshold mechanosensitive cutaneous afferent (all soft-brush sensitive) was classified by its physiological characteristics, as per the criteria used in [44], [71]. Briefly, individual Aβ low-threshold mechanoreceptors were separated into rapidly and slowly adapting types based on their adaptive responses to ramp-and-hold indentation of the skin. Three groups of rapidly adapting units were identified as follows: HFA, responsive to hair deflection and light air puffs; FA-II, comprising a single spot of maximal sensitivity and robust response to remote tapping; Field, comprising multiple spots of high sensitivity with no response to hair displacement or remote tapping of the skin. Two groups of slowly adapting (types I and II) were identified where several features were examined including spontaneous firing, stretch sensitivity, and receptive field characteristics. In addition, an inter-spike interval pattern to sustained indentation (100 mN for 30 s) was tested. Coefficients of variation of inter-spike intervals for all SA-IIs (n=4) were in the range of 0.15 to 0.23. This was also measured for one SA-I and its coefficient of variation was 1.92. These values are consistent with previous observations [24], [44], [72]. Single muscle spindle afferents were identified by stretch of the receptor-bearing muscle along its line of action. These were not further classified into primary and secondary afferents.

Mechanical thresholds of all cutaneous afferent fibers were measured using Semmes-Weinstein monofilaments (nylon fiber; Aesthesio, Bioseb, Pinellas Park, FL, USA), except HFA whose preferred stimulus is hair movement so responses to light air puffs were determined. The monofilaments were applied manually with a rapid onset until the monofilament buckled: If a unit responded to the same (weakest) monofilament in at least 50% of trials, it was taken as the mechanical threshold. Based on prior work showing that 4 mN threshold divides the low threshold (<4 mN) and high threshold (≥ 4 mN) cutaneous afferent populations in hairy skin [24], [71], only those afferents with thresholds below 4 mN were considered. Further, any cutaneous afferent with a receptive field located at a site inaccessible for the delivery of expressions was discarded.

All neural data were recorded and processed using LabChart Pro for setup 1 (v8.1.5 and PowerLab 16/35 hardware PL3516/P, ADInstruments, Oxford, UK) and SC/ZOOM for setup 2 (Physiology Section, Department of Integrative Medical Biology, Umeå University). Action potentials were distinguished from background noise with a signal-to-noise ratio of at least 2:1 and were confirmed to have originated from the recorded afferent by a semi-automatic inspection of their morphology. For further details see [24].

### Statistical and classification analyses

When examining the basic firing properties of the afferents (Fig. 2), per expression trial, the mean IFF was calculated over the whole 10 s neural recording and the number of spikes was calculated from a 1 s chunk constraining the largest number of spikes. The duration of 1 s was determined as it could be covered by non-stop contact across all expressions. Mann-Whitney U tests were conducted for pairwise comparisons of those two properties across afferent subtypes and touch gestures. Post-hoc Benjamini-Hochberg method was used for multiple testing correction.

A five-layer 1D CNN model and an SVM model were employed for classification analysis. The structure and hyper parameters of the CNN model were determined by cross validation grid search with data from all afferent subtypes combined together and there were 16,646 trainable parameters in total. For each layer of CNN, 0.2 dropout was applied. The model was trained with Early Stopping and the ADAM optimizer with a reducing learning rate starting from 0.001. The same model was trained and tested for each afferent subtype separately based on the loss of categorical cross-entropy. For classification using full 10 s recordings and the identified best segments, five-fold cross validation were repeated 20 times to obtain the average prediction results for both CNN and SVM models (Fig. 3, 5B, 5C). Among five neural firing properties extracted for SVM classification (Fig. 3B, 5B, 5C, S1B), the number of spikes was calculated from full 10 s, IFF variation was calculated as the coefficient of variation of IFF, and the number of bursts was defined as the number of spike bursts separated by gaps of inter-spike intervals larger than 1 s. For the identification of the most informative segments of neural recordings (Fig. 4), window position metrics of IFF variation and IFF entropy were calculated from step-interpolated IFF to better reflect the time-series pattern of touch expressions. The sampling rates of window length were designed as every 0.1 s from 0.1 s to 4 s, and every 0.25 s from 4 s to 10 s. Five-fold cross validation of CNN was repeated twice across all window lengths to identity the best window position metrics, where Mann-Whitney U tests and post-hoc Benjamini-Hochberg correction were applied for pairwise comparison (Fig. 4B). Best window lengths were identified based on seven repeats of five-fold cross validation of CNN across all window lengths and the best two window position metrics (Fig. 4D, 5E). For the spike-time sensitivity analysis, average accuracies were obtained from five repeats of five-fold cross validation with each level of noise tested by ten different sets of random noise.

