## Supplemental information for "Mechanoreceptive Aβ primary afferents discriminate naturalistic social touch inputs at a functionally relevant time scale"

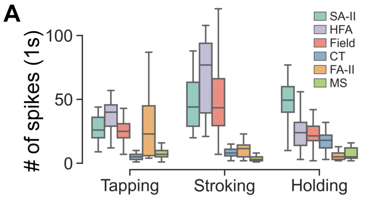


| Tapping | SA-II | HFA | Field | CT | FA-II | MS |
|---|---|---|---|---|---|---|
| SA-II | | *** | - | **** | - | **** |
| HFA | | | *** | **** | * | **** |
| Field | | | | **** | - | **** |
| CT | | | | | *** | * |
| FA-II | | | | | | *** |
| MS | | | | | | |

| Stroking | SA-II | HFA | Field | CT | FA-II | MS |
|---|---|---|---|---|---|---|
| SA-II | | **** | - | **** | **** | **** |
| HFA | | | * | **** | **** | **** |
| Field | | | | **** | **** | **** |
| CT | | | | | - | **** |
| FA-II | | | | | | **** |
| MS | | | | | | |

| Holding | SA-II | HFA | Field | CT | FA-II | MS |
|---|---|---|---|---|---|---|
| SA-II | | *** | **** | **** | **** | **** |
| HFA | | | - | - | *** | **** |
| Field | | | | - | *** | **** |
| CT | | | | | ** | *** |
| FA-II | | | | | | - |
| MS | | | | | | |

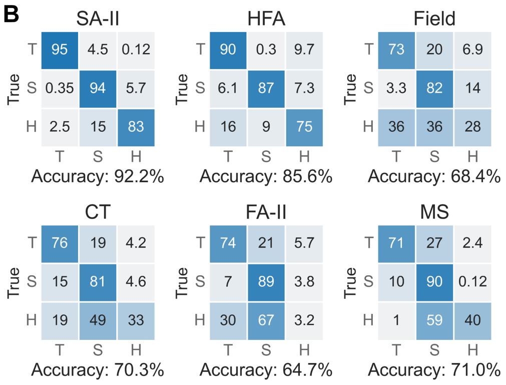


*Figure S1.* (A) Comparison of the number of spikes across afferent subtypes per gesture. *$p < 0.05$, **$p < 0.01$, ***$p < 0.001$, ****$p < 0.0001$ were derived by Mann–Whitney U tests with Benjamini-Hochberg post-hoc correction. (B) SVM classification of touch gestures per afferent subtype using five firing properties the same as Fig. 2B. “T” represents tapping, “S” represents stroking, and “H” represents holding.

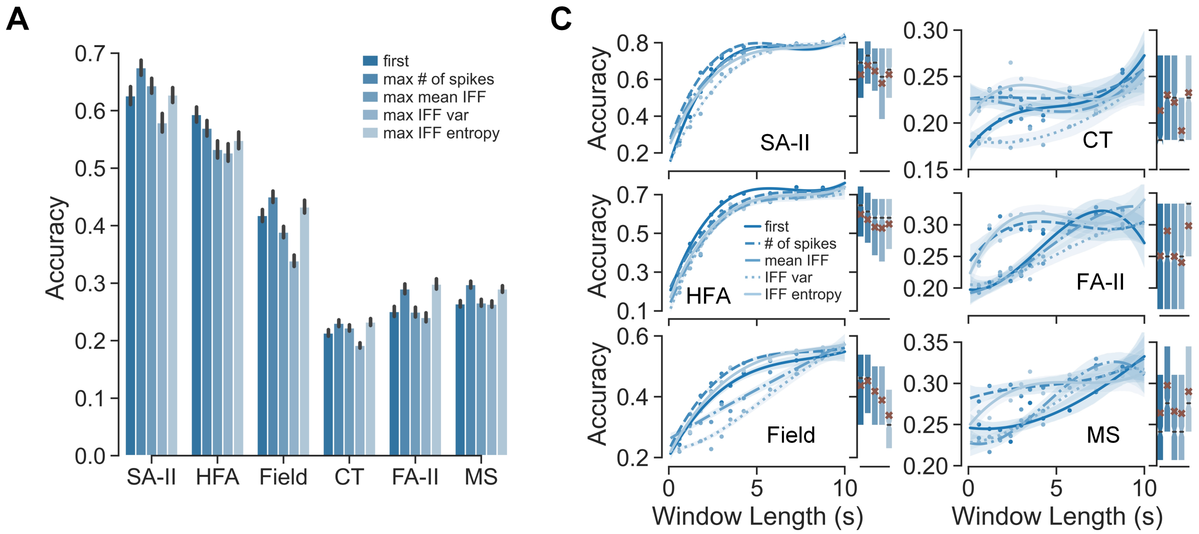


B

| SA-II **** | first | count | mean | var | entropy |
|---|---|---|---|---|---|
| first | 0.63 | *** | - | **** | - |
| count | | 0.67 | *** | **** | **** |
| mean | | | 0.64 | **** | - |
| var | | | | 0.58 | *** |
| entropy | | | | | 0.63 |

| HFA **** | first | count | mean | var | entropy |
|---|---|---|---|---|---|
| first | 0.59 | ** | **** | **** | **** |
| count | | 0.57 | ** | *** | - |
| mean | | | 0.53 | - | - |
| var | | | | 0.53 | - |
| entropy | | | | | 0.55 |

| Field **** | first | count | mean | var | entropy |
|---|---|---|---|---|---|
| first | 0.42 | **** | *** | **** | - |
| count | | 0.45 | **** | **** | * |
| mean | | | 0.39 | **** | **** |
| var | | | | 0.34 | **** |
| entropy | | | | | 0.43 |

| CT **** | first | count | mean | var | entropy |
|---|---|---|---|---|---|
| first | 0.21 | *** | * | **** | **** |
| count | | 0.23 | - | **** | - |
| mean | | | 0.22 | **** | - |
| var | | | | 0.19 | **** |
| entropy | | | | | 0.23 |

| FA-II **** | first | count | mean | var | entropy |
|---|---|---|---|---|---|
| first | 0.25 | **** | - | - | **** |
| count | | 0.29 | **** | **** | - |
| mean | | | 0.25 | - | **** |
| var | | | | 0.24 | **** |
| entropy | | | | | 0.30 |

| MS **** | first | count | mean | var | entropy |
|---|---|---|---|---|---|
| first | 0.26 | **** | - | - | **** |
| # | | 0.30 | **** | **** | - |
| mean | | | 0.27 | - | **** |
| var | | | | 0.26 | **** |
| entropy | | | | | 0.29 |

*Figure S2.* (A) Classification accuracies across window position metrics averaged over all window lengths for each afferent subtype. (B) *$p < 0.05$, **$p < 0.01$, ***$p < 0.001$, ****$p < 0.0001$ were derived by Mann–Whitney U tests with Benjamini-Hochberg post-hoc correction. (C) Classification accuracies across window position metrics along with the increase of window length. Curves were fitted using third-order polynomial functions, points denote means of 10 evenly-binned data. Bar plots show distributions of classification accuracies over all window lengths per window position metric, and brown cross markers denote means per window position metric.

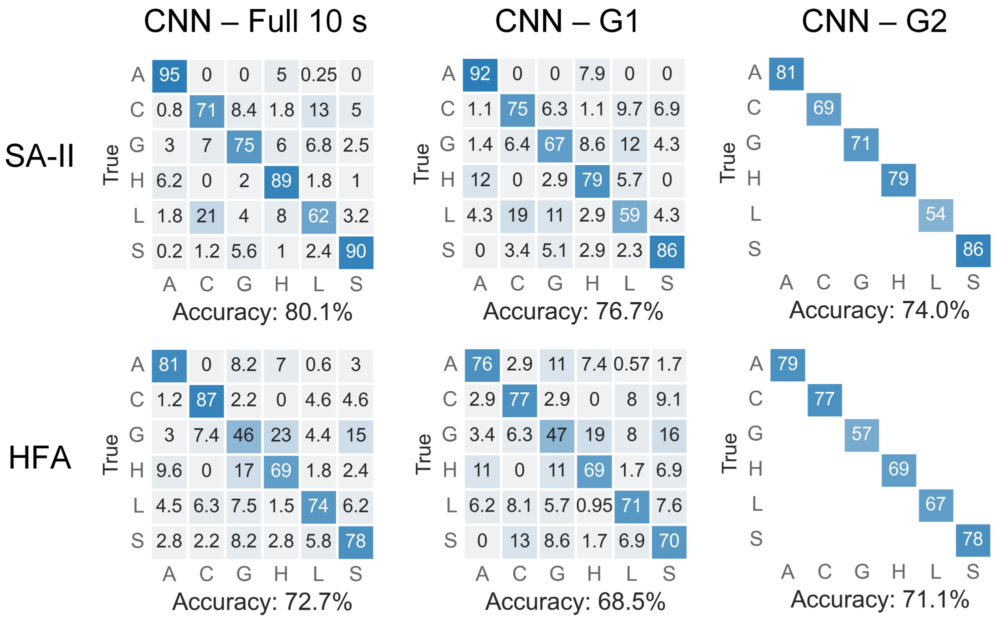


*Figure S3.* Classification results using CNN. G1, G2 refer to two groups of neural recording chunks derived by saturation window lengths per afferent subtype (Fig. 4D) and saturation window lengths per afferent-expression combination, respectively (Fig. 5D). As CNN requires the same length of input data, for G2, classification of each expression was conducted using its corresponding saturation window length for all neural recordings. Only the accuracy for that expression was shown in the confusion matrix, accuracies for other expression were left as blank.

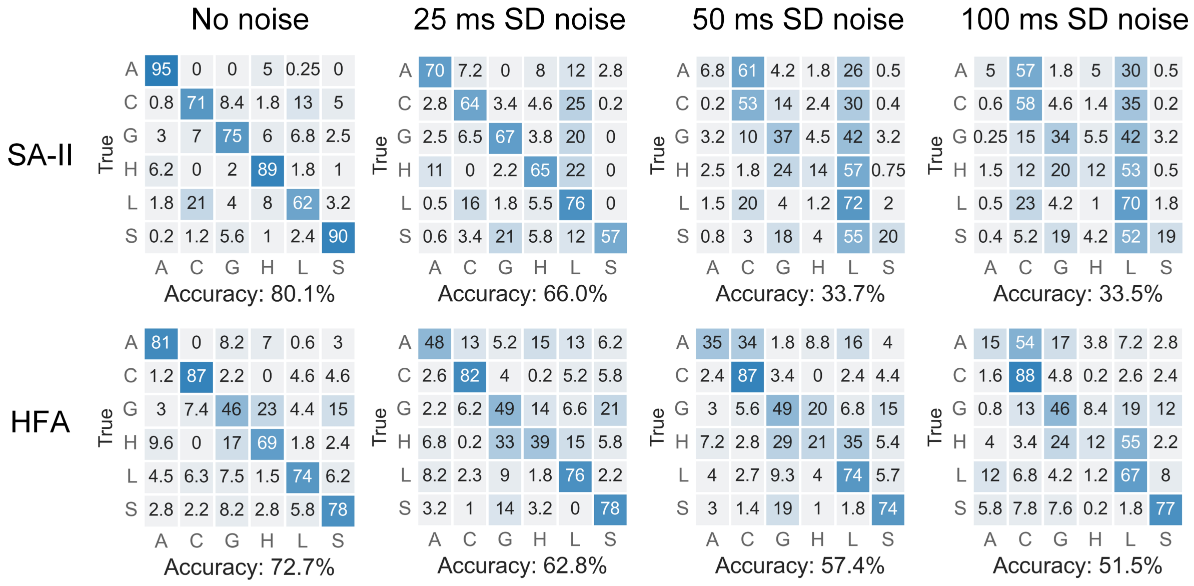


*Figure S4.* CNN classification accuracies using 10 s neural recordings with different level of noise added.

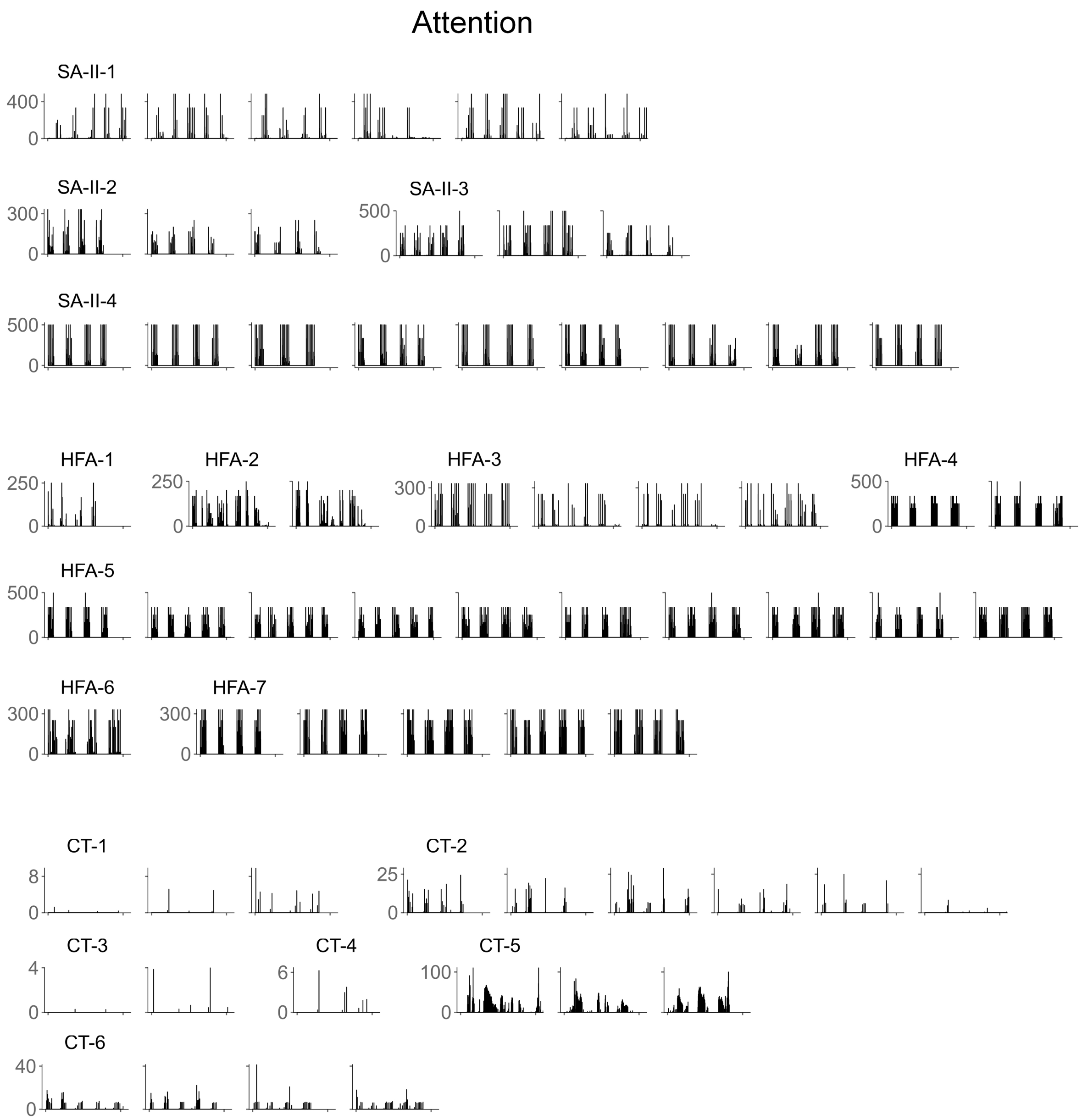


*Figure S5.* Neural recordings of SA-II, HFA, CT subtypes for the attention expression. All trials are shown here in the format of Instantaneous firing frequency (IFF, Hz).
